# Rethinking Margin of Stability: Incorporating Step-To-Step Regulation to Resolve the Paradox

**DOI:** 10.1101/2021.12.01.470263

**Authors:** Meghan Kazanski, Joseph P. Cusumano, Jonathan B. Dingwell

## Abstract

Derived from inverted pendulum dynamics, mediolateral Margin of Stability (*MoS*_*ML*_) is a mechanically-grounded measure of instantaneous stability. However, average *MoS*_*ML*_ measures yield paradoxical results. Gait pathologies or perturbations often induce larger (supposedly “more stable”) average *MoS*_*ML*_, despite clearly *de*stabilizing factors. However, people do not walk “on average” – they walk (and sometimes lose balance) one step at a time. We assert the paradox arises because averaging discards step-to-step dynamics. We present a framework unifying the inverted pendulum with Goal-Equivalent Manifold (GEM) analyses. We identify in the pendulum’s center-of-mass dynamics constant-*MoS*_*ML*_ manifolds, including one candidate “stability GEM” signifying the goal to maintain some constant 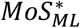. We used this framework to assess step-to-step *MoS*_*ML*_ dynamics of humans walking in destabilizing environments. While goal-relevant deviations were readily corrected, humans did not exploit equifinality by allowing deviations to persist *along* this GEM. Thus, maintaining a constant 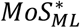 is inconsistent with observed step-to-step fluctuations in center-of-mass states. Conversely, the extent to which participants regulated fluctuations in foot placements strongly predicted regulation of center-of-mass fluctuations. Thus, center-of-mass dynamics may arise *in*directly as a consequence of regulating mediolateral foot placements. To resolve the paradox caused by averaging *MoS*_*ML*_, we present a new statistic, Probability of Instability (*PoI*_*L*_), to predict instability likelihood. Participants exhibited increased *PoI*_*L*_ when *de*stabilized (p = 9.45×10^−34^), despite exhibiting larger (“more stable”) average *MoS*_*ML*_ (p = 1.70×10^−15^). Thus, *PoI*_*L*_ correctly captured people’s increased risk of losing lateral balance, whereas average *MoS*_*ML*_ did not. *PoI*_*L*_ also explains why peoples’ average *MoS*_*ML*_ increased in destabilizing contexts.

## INTRODUCTION

Falls are a leading cause of injury among young (Cho et al., 2021), middle-age (Wang et al., 2021) and older (Burns and Kakara, 2018) adults. Most people fall while walking (Heijnen and Rietdyk, 2016) and walking humans are intrinsically less stable laterally (Bauby and Kuo, 2000; McAndrew Young et al., 2012). Consequently, sideways falls are particularly common (Crenshaw et al., 2017) and injurious (Yang et al., 2020) in older adults. Generally, humans can prevent or overcome mediolateral instability by adjusting foot placements (Townsend, 1985) to redirect their center-of-mass (CoM) dynamics (Bruijn and van Dieën, 2018). However, we still lack a comprehensive framework to describe how humans regulate their movements *from step-to-step* to maintain mediolateral stability.

The inverted-pendulum model of frontal-plane dynamics represents the simplest conceptualization of the mediolateral balance problem during walking (Fig. 1A). Derived from this model, the mediolateral Margin of Stability, *MoS*_*ML*_ (Hof et al., 2005), provides a mechanically-grounded measure (as a physical distance) of how close this system is to becoming laterally unstable at any instant. Hof’s condition for dynamic stability is met when the vertical projection of the ‘extrapolated’ CoM remains within the base-of support boundaries (Hof et al., 2005). In the lateral direction, any value of 0 ≤ *MoS*_*ML*_ ≤ *w*_*foot*_ meets this condition, with larger values interpreted to indicate greater lateral stability. Dynamics that exhibit *MoS*_*ML*_ outside this inequality *might* result in a sideways fall, unless Hof’s condition is restored via active rebalancing mechanisms not incorporated into the inverted pendulum model, like taking a step. *MoS*_*ML*_ thus approximates the biomechanical requirement for maintaining lateral balance *within* any step. However, Hof’s condition must then be re-established *at each new step*.

**Figure 1.**
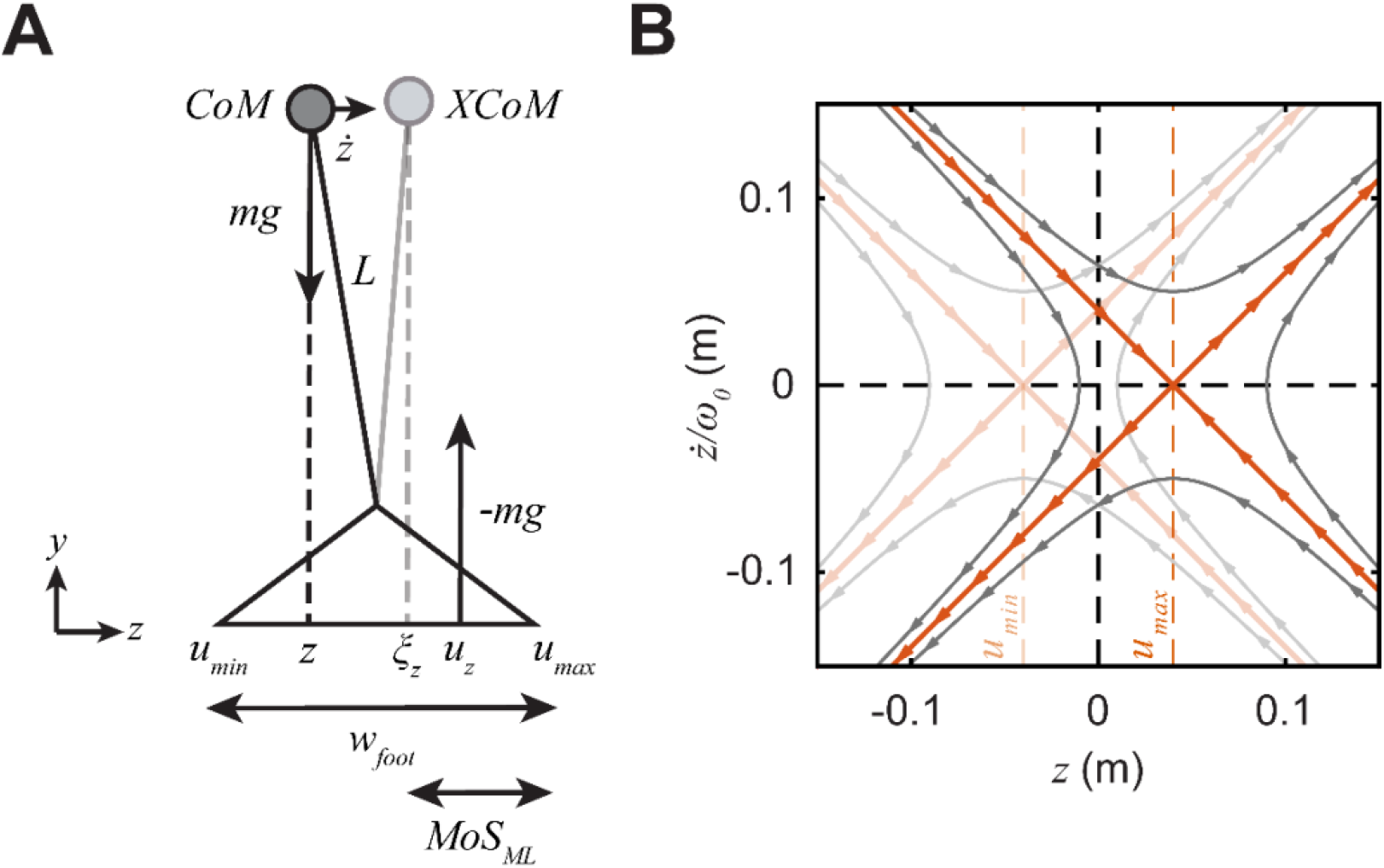
**(A)** The frontal plane inverted pendulum exemplifies mediolateral CoM dynamics (*z, ż*) during single-stance of each step (adapted from (Hof et al., 2005)). Motion is modeled as that of a point mass *m* located at the body CoM, attached a distance *L* from the ankle joint of the finite-width foot (base-of-support; *w*_*foot*_). The mediolateral center of pressure, *u*_*z*_, can act at any of infinite possible locations between *u*_*min*_ ≤ *u*_*z*_ ≤ *u*_*max*_. Hof’s condition for dynamic stability is met when the vertical projection of the “extrapolated” CoM (*ξ*_*z*_ = *z* + *ż/ω*_*0*_) lies within this same range: i.e., *u*_*min*_ ≤ *ξ*_*z*_ ≤ *u*_*max*_. The mediolateral margin of stability (*MoS*_*ML*_; Eq. (1)) is the medial-to-lateral distance from *ξ*_*z*_ to *u*_*max*_ (lateral foot boundary). A corresponding lateral-to-medial margin of stability (not shown) also exists as the distance from *ξ*_*z*_ to *u*_*min*_. **(B)** The CoM state space of the inverted pendulum (adapted from (Bottaro et al., 2008; Hof, 2008)) for different constant values of *u*_*z*_. The horizontal axis (*z*) is the mediolateral CoM position, where *z* = 0 denotes the center of the foot. The vertical axis (*ż/*ω_*0*_) is the scaled mediolateral CoM velocity. Example “snap-shots” of topologically-equivalent phase portraits are plotted for the two outermost cases *u*_*z*_ = *u*_*max*_ (dark lines) and *u*_*z*_ = *u*_*min*_ (light lines). Thick orange lines are the stable manifolds (negative diagonals) and unstable manifolds (positive diagonals) of each saddle. Grey flow lines direct the passive CoM dynamics during a step (flow direction indicated by arrows). There exists an infinite set of such phase portraits, and thus an infinite set of such stable manifolds, for all possible *u*_*z*_ between *u*_*min*_ ≤ *u*_*z*_ ≤ *u*_*max*_.

One might also expect *MoS*_*ML*_ to reflect falling likelihood. However, applying *MoS*_*ML*_ to walking produces counterintuitive results when values are averaged across steps. People exhibit larger (i.e., supposedly more stable) average *MoS*_*ML*_ when subjected to *de-*stabilizing perturbations (Beltran et al., 2014; McAndrew Young et al., 2012; Onushko et al., 2019). Likewise, multiple fall-prone populations exhibit larger average *MoS*_*ML*_ (Watson et al., 2021), including older adults (Hurt and Grabiner, 2015) and persons with stroke (Tisserand et al., 2018), amputation (Gates et al., 2013; Hof et al., 2007; Rodrigues et al., 2021), cerebral palsy (Rethwilm et al., 2021), spinal cord injury (Day et al., 2012), Parkinson’s disease (Lencioni et al., 2021) or multiple sclerosis (Peebles et al., 2016). Paradoxically then, despite clearly destabilizing factors, these individuals’ average *MoS*_*ML*_ should classify them as *more* stable (not less) according to Hof’s definition.

We suggest this paradox arises largely from averaging *MoS*_*ML*_ values across multiple steps. People do not fall “on average” – they fall from singular events (e.g., perturbations) that occur in real time (i.e., during a step). Averaging *MoS*_*ML*_ inherently discards any information about how *MoS*_*ML*_ varies across steps and/or what people might adjust to maintain or recover balance. People likely adjust foot placement to modulate mediolateral balance (Bruijn and van Dieën, 2018). CoM state deviations predict *current* step foot placements (Patil et al., 2019; Wang and Srinivasan, 2014) – and those foot placements then redirect the CoM on the *next* step (Redfern and Schumann, 1994; Townsend, 1985). We must therefore examine how humans *regulate* relevant gait variables *across steps* to maintain mediolateral stability. Here, “regulation” denotes the corrective processes people enact from step to step to achieve specific goal-directed locomotor tasks (Patil et al., 2022).

The Goal-Equivalent Manifold (GEM) concept (Cusumano and Cesari, 2006) allows us to formulate task goal functions and control models to pose testable hypotheses regarding how humans regulate stepping to walk (Dingwell et al., 2010). If we hypothesize a particular walking task goal, the GEM defines the set of all possible solutions that equally satisfy that goal. We can then test this hypothesis by quantifying how relevant stepping variables fluctuate about this GEM from step to step. We expect “GEM-aware” walking dynamics to generally follow the Minimal Intervention Principle (MIP; (Todorov, 2004)), where error correction processes strongly drive goal-relevant deviations onto the GEM, but only weakly adjust goal-equivalent deviations along it (Cusumano and Dingwell, 2013; Dingwell et al., 2010). The GEM framework successfully describes how humans regulate mediolateral foot placement during unperturbed (Dingwell and Cusumano, 2019) and perturbed (Dingwell et al., 2021; Kazanski et al., 2020) walking, or when given explicit task feedback (Render et al., 2021). We expect the same GEM concepts should be useful for studying how people regulate their movements to maintain mediolateral stability.

Here, we first present a novel analytical framework that unifies the inverted pendulum model with GEM-based analyses to re-conceptualize the mediolateral balance problem. We identify in the inverted pendulum’s CoM dynamics an infinite set of constant-*MoS*_*ML*_ manifolds. Prior experiments led others to speculate that people may try to maintain some constant *MoS*_*ML*_ as they walk (Curtze et al., 2011; Hof, 2008; Rosenblatt and Grabiner, 2010). This suggests that some particular constant-*MoS*_*ML*_ manifold in the inverted pendulum’s CoM dynamics might serve as a candidate “stability GEM”, reflecting a walker’s goal to maintain that *MoS*_*ML*_ value from step to step. Here, we directly test this hypothesis by quantifying the step-to-step fluctuation dynamics of minimum *MoS*_*ML*_ time series about the proposed candidate stability GEM. We test how these fluctuation dynamics vary with age and externally-imposed perturbations.

Conversely, because the CoM is not located at any particular point in the body (and need not even reside *within* the body), humans arguably can neither directly perceive (Ting and Macpherson, 2004) nor actuate their CoM itself. Thus, it is not *a priori* obvious walkers can directly correct CoM deviations from any constant value of *MoS*_*ML*_ taken as a walking task goal. Instead, observed CoM dynamics may arise from other directly regulated processes (like foot placement, etc.) that *in*directly adjust CoM states (Hof, 2007; Li and Huang, 2022). We therefore tested the alternative hypothesis that how people regulate mediolateral foot placement fluctuations (Kazanski et al., 2020) statistically explains mediolateral CoM state fluctuations. If confirmed, this would suggest that the structure of step-to-step fluctuations in people’s mediolateral CoM dynamics arise as a consequence of how they regulate mediolateral foot placements.

Lastly, if people do regulate their movements from step to step to achieve specific margin of stability goals, this likely reflects the extent to which they consider instability *risk* (i.e., the likelihood of experiencing an adverse event). In reaching tasks, humans use “risk-sensitive” control strategies (Braun et al., 2011) that seek to minimize variance *per se* (John et al., 2016; Nagengast et al., 2011). People may implement similar strategies during walking: e.g., to trade off mediolateral stability for effort (Dean et al., 2007) and/or maneuverability (Hsieh et al., 2018). If humans seek to mitigate risk of instability, the step-to-step variance of mediolateral CoM dynamics should be structured accordingly. However, neither instantaneous nor average measures of *MoS*_*ML*_ capture this. Here, we propose a new *MoS*_*ML*_-based measure, the Probability of Instability (*PoI*), that directly predicts mediolateral instability likelihood on any *given* step. We demonstrate that this new metric resolves the long-standing Margin of Stability paradox.

## METHODS

### Participants, Protocol, and Data Processing

We conducted the analyses presented here on data from a cohort of young (YH) and older (OH) healthy adults previously described in Kazanski et al. (2020) (Table 1). Complete descriptions of participants, assessments, experimental protocol, and data processing methods are in *Supplement 1*.

**Table 1.** Relevant participant characteristics for Young (YH) and Older (OH) healthy adults. All values except Sex are given as mean ± standard deviation. Two-sample t-test results (p-values) for Age group differences are shown. For additional details, see Supplement S1 and Kazanski et al. (2020).

| Characteristic: | Young Healthy (YH): | Older Healthy (OH): | p-value |
| --- | --- | --- | --- |
| Sex [M/F] | 8 / 9 | 7 / 10 | N/A |
| Age [yrs] | 23.7 ± 3.7 | 67.5 ± 4.9 | < **0.001** |
| Body Height [m] | 1.72 ± 0.1 | 1.71 ± 0.1 | 0.772 |
| Body Mass [kg] | 64.5 ± 12.5 | 73.3 ± 18.9 | 0.118 |
| Body Mass Index [kg/m<sup>2</sup>] | 21.7 ± 3.2 | 24.9 ± 5.4 | **0.042** |
| Leg Length [m] | 0.89 ± 0.05 | 0.90 ± 0.05 | 0.453 |
| **Assessment:** |  |  |  |
| Timed Up and Go [s] | 8.06 ± 1.20 | 8.66 ± 1.08 | 0.139 |
| Four Square Step Test [s] | 7.18 ± 1.53 | 9.07 ± 2.07 | **0.005** |
| Preferred Walking Speed [m/s] | 1.46 ± 0.15 | 1.35 ± 0.14 | **0.033** |

Briefly, each participant walked in a “V-Gait” virtual reality system, (Motekforce Link, Amsterdam, Netherlands) in each of 3 conditions: normal walking with no perturbations (NOP), or walking with continuous mediolateral oscillations of either the visual field (VIS) or treadmill platform (PLAT). We analyzed kinematic data from markers placed on each heel, lateral malleolus, and pelvis (Havens et al., 2018). We extracted stepping data (Zeni et al., 2008) for *N*=230 consecutive steps for each trial.

### Mediolateral Margin of Stability

We defined the *z-*coordinate as mediolateral (Fig. 1A) and used pelvic centroid motion to approximate the continuous mediolateral CoM state (*z, ż*). We used the mediolateral position of the leading foot’s lateral malleolus marker to define the lateral base-of-support boundary (*u*_*max*_). Using Hof’s expression (Hof et al., 2005), we computed the *minimum* mediolateral Margin of Stability (*MoS*_*ML*_) for each step *n* as:

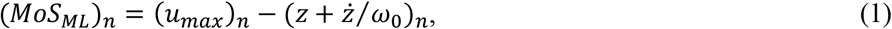

where (*z*+*ż/ω*_0_) is the mediolateral extrapolated CoM position accounting for CoM velocity, normalized by the eigenfrequency 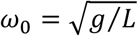 of a pendulum of length *L* = 1.34×*l*, where *l* is leg length measured from the lateral malleolus to greater trochanter.

For each trial, we extracted time series of *MoS*_*ML*_ for all *n* ∈ (1,…, *N*) steps. We computed the mean and standard deviation (*σ*) of each *MoS*_*ML*_ time series. We used Detrended Fluctuation Analysis (DFA) to compute scaling exponents (α) quantifying the degree of statistical persistence in each time series (Maraun et al., 2004): values of α > 0.5 indicate statistical persistence, α = 0.5 indicates uncorrelated deviations, and α < 0.5 indicates anti-persistence. Following the GEM/MIP theoretical framework (Dingwell and Cusumano, 2010; Dingwell et al., 2010), time series exhibiting α ≈ 0.5 typify strongly-regulated variables with rapid step-to-step corrections. Conversely, time series exhibiting α >> 0.5 typify weakly-regulated variables with deviations that go uncorrected over many steps. For each dependent measure, we performed a two-factor mixed-effects (Age×Condition) analysis of variance (ANOVA) (*Supplement 2*).

### Candidate Stability Goal-Equivalent Manifold

*MoS*_*ML*_ (Eq. (1)) assumes mediolateral balance is well-approximated by an inverted pendulum with finite base-of-support with bounds [*u*_*min*_, *u*_*max*_] (Fig. 1A). For any fixed value of center-of-pressure (*u*_*z*_), these dynamics define a single phase portrait in [*z, ż/ω*_0_] (Bottaro et al., 2008; Hof, 2008). Across *all* possible values of *u*_*min*_ ≤ *u*_*z*_ ≤ *u*_*max*_, one can then heuristically conceive of an infinite *family* of such phase portraits (Fig. 1B). So long as the extrapolated CoM (*ξ*_*z*_) remains within *u*_*min*_ ≤ *ξ*_*z*_ ≤ *u*_*max*_, the walker can modulate their center-of-pressure within these bounds to maintain balance (Hof et al., 2005).

Relating these pendulum dynamics to Hof’s *Margin* of Stability, we rearrange Eq. (1) as:

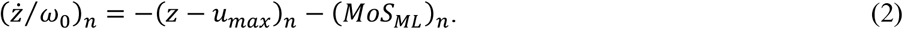

Now, in the [*z-u*_*max*_, *ż/ω*_0_] plane (Fig. 2A), Eq. (2) specifies a diagonal line (slope −1; intercept −*MoS*_*ML*_) that defines a *constant-MoS*_*ML*_ *manifold*: along any such diagonal, infinite combinations of [(*z-u*_*max*_)_*n*_, (*ż/ω*_0_)_*n*_] yield the exact same *MoS*_*ML*_. In particular, the two diagonals at *MoS*_*ML*_ = 0 and *MoS*_*ML*_ = *w*_*foot*_ in Fig. 2A correspond to the stable manifolds of the two phase portraits at *u*_*z*_ = *u*_*max*_ and *u*_*z*_ = *u*_*min*_ in Fig. 1B, respectively. These boundaries form a “Stability” region, outside of which a walker fails to meet Hof’s condition (Fig. 2A).

**Figure 2.**
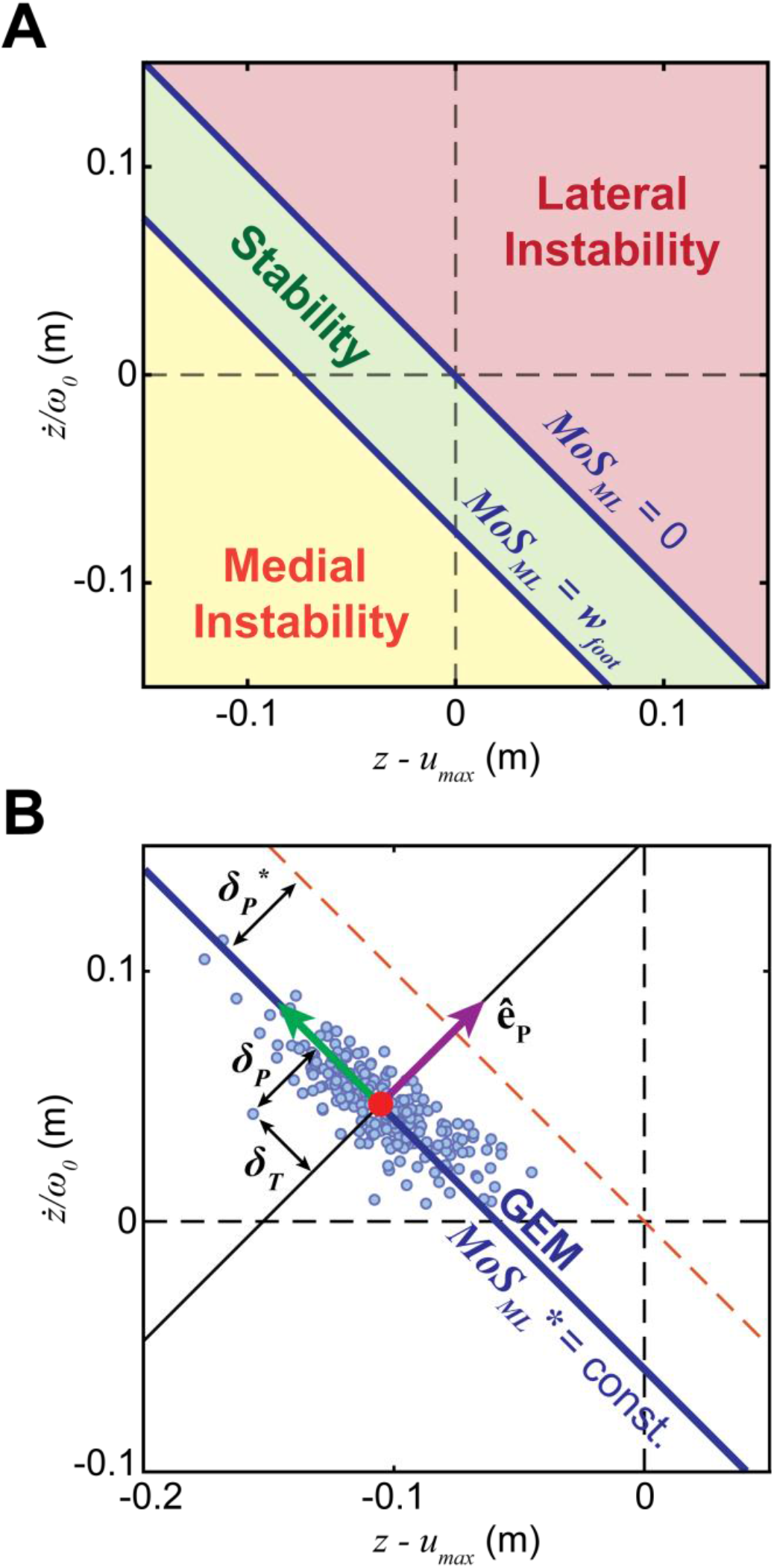
**(A):** From Hof’s (Hof et al., 2005) definition of mediolateral Margin of Stability (*MoS*_*ML*_; Eq. (1)), level sets of constant *MoS*_*ML*_ define *constant-MoS*_*ML*_ *manifolds* as diagonal lines in the [*z-u*_*max*_, *ż/ω*_0_] plane (Eq. (2)). Infinite such manifolds exist for all possible values of *MoS*_*ML*_. Incorporating Hof’s condition for dynamic stability (i.e., 0 ≤ *MoS*_*ML*_ ≤ *w*_*foot*_; Eq. (3)), these level sets then partition the [*z*-*u*_*max*_, *ż*/ω_0_] plane into three distinct regions. The *MoS*_*ML*_ = 0 manifold defines the boundary between Stability (green) and Lateral Instability (red) regions. The *MoS*_*ML*_ = *w*_*foot*_ manifold defines the boundary between the Stability and Medial Instability (yellow) regions. **(B):** One such constant-*MoS*_*ML*_ manifold (diagonal blue line) depicted as a candidate *goal-equivalent manifold* (GEM) in [*z*-*u*_*max*_, *ż*/ω_0_] to achieve some desired constant 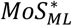 (Eq. 4). Data points (filled blue markers) represent single steps from a typical trial, taken at each minimum *MoS*_*ML*_. Unit vectors [*ê*_*T*_, *ê*_*P*_] define directions tangent and perpendicular to this candidate GEM from the mean operating point (red filled circle). Time series of CoM states are transformed into deviations tangent (*δ*_*T*_) and perpendicular (*δ*_*P*_) to the GEM (Eq. 5). *δ*_*P*_^*^ is the distance along *ê*_*P*_ from the 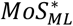 GEM to the *MoS*_*ML*_ = 0 lateral stability boundary.

Hof’s condition for dynamic stability thus suggests a possible task *goal* for walking, namely to achieve *MoS*_*ML*_ values within this Stability region for all *n* steps in a sequence of *N* consecutive steps:

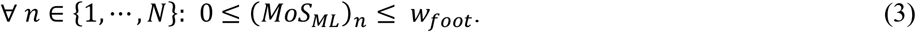

Because *any* sequence of [(*z-u*_*max*_)_*n*_, (*ż/ω*_0_)_*n*_] that satisfies Eq. (3) – and there are *infinite* such combinations – meets this condition, many strategies exist to generate such feasible stepping sequences. If, as suggested (Curtze et al., 2011; Hof, 2008; Rosenblatt and Grabiner, 2010), people try to maintain some constant *MoS*_*ML*_ as they walk, this would mean they try to satisfy Eq. (3) by maintaining some constant 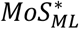, such that 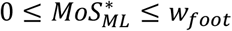, at each step. We can now express this candidate stabilizing approach using the *goal function*:

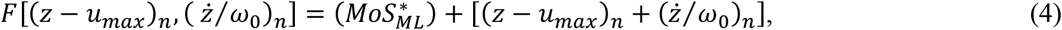

where the person’s goal is to drive *F* → 0 at each step. Eq. (4) thus defines a candidate *Goal-Equivalent* Manifold (GEM) for mediolateral stability as a specific constant-*MoS*_*ML*_ manifold achieving 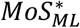 (Fig. 2B). Here, we test the hypothesis that humans adopt such a constant-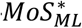approach to regulate walking.

For each trial, we defined the candidate GEM by 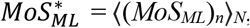, where ⟨•⟩_*N*_ denotes the average over *N* steps. Following (Dingwell et al., 2010), we defined an “operating point” on the GEM as (*z-u*_*max*_)^*^ = ⟨(*z-u*_*max*_)_*n*_⟩_*N*_ and (*ż/ω*_0_)^*^ = ⟨(*ż/ω*_0_)_*n*_⟩_*N*_. We defined deviations in [(*z-u*_*max*_)_*n*_, (*ż/ω*_0_)_*n*_] from this operating point as (*z-u*_*max*_)’_*n*_ = (*z-u*_*max*_)_*n*_ *−* (*z-u*_*max*_)^*\**^ and (*ż/ω*_0_)’_*n*_ = (*ż/ω*_0_)_*n*_ probability density function of a given time series *−* (*ż/ω*_0_)_*n*_^*\**^. We then linearly transformed [(*z-u*_*max*_)’_*n*_, (*ż/ω*_0_)’_*n*_] coordinates into [(*δ*_*T*_)_*n*_, (*δ*_*P*_)_*n*_] coordinates along GEM-specific unit vectors [*ê*_*T*_, *ê*_*P*_] (Fig. 2B):

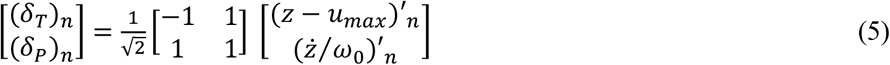

Deviations perpendicular to the GEM (*δ*_*P*_) are goal-*relevant*: they directly impact *MoS*_*ML*_. Conversely, deviations tangent to the GEM (*δ*_*T*_) are goal-*equivalent*: they have no impact on *MoS*_*ML*_. Walking humans readily correct, from step-to-step, goal-relevant deviations perpendicular to GEMs that reflect identified walking goals (Dingwell et al., 2010). Conversely, they allow goal-equivalent deviations to persist (i.e., go uncorrected) across many steps, exploiting redundancy consistent with the Minimal Intervention Principle (Todorov, 2004).

We computed time series of *δ*_*P*_ and *δ*_*T*_ for each trial. For each, we computed variability (*σ*) to quantify average deviation magnitude and statistical persistence (DFA *α*) to quantify strength of step-to-step corrections. If participants exploited the available redundancy along the candidate stability GEM, we predicted they would: (1) reduce goal-relevant deviations from the GEM (i.e., *σ*(*δ*_*P*_) << *σ*(*δ*_*T*_)) and (2) readily correct goal-relevant deviations (i.e., α(*δ*_*P*_) ≈ 0.5), while also allowing goal-equivalent deviations along the GEM to persist (i.e., α(*δ*_*T*_) >> 0.5) (Dingwell et al., 2010). For each dependent measure, we conducted three-factor and two-factor mixed-effects ANOVAs to test these predictions (see *Supplement 2*).

### Regulating Foot Placement to Maintain Mediolateral Balance

Humans likely cannot perceive their CoM directly (Ting and Macpherson, 2004), nor can they directly actuate it. Instead, step-to-step fluctuations in CoM state [(*z-u*_*max*_)_*n*_, (*ż/ω*_0_)_*n*_] could arise from other processes that redirect the CoM between steps. Foot placement is arguably the main such process, as walking humans systematically regulate step-to-step lateral foot placements (Dingwell and Cusumano, 2019), especially when balance is challenged (Dingwell et al., 2021; Kazanski et al., 2020).

We therefore tested an alternative hypothesis that step-to-step CoM dynamics might instead arise as a consequence of how participants regulated step-to-step fluctuations in step width (*w*) and lateral body position (*z*_*B*_), which we previously reported in (Kazanski et al., 2020). Here, we computed pairwise (trial-by-trial) Pearson correlations between our previous measures of α(*w*) and α(*z*_*B*_) and currently calculated measures of α(*δ*_*T*_) and α(*δ*_*P*_). These correlations assessed whether, across age groups and conditions, the extent to which participants corrected step-to-step deviations in mediolateral foot placement (α(*w*), α(*z*_*B*_)) predicted the corresponding fluctuation properties (α(*δ*_*T*_), α(*δ*_*P*_)) in CoM state.

### Probability of Instability (PoI)

It is evident from Fig. 2 that average values of *MoS*_*ML*_ cannot predict lateral instability likelihood. While *MoS*_*ML*_ is a meaningful measure of one’s instantaneous stability state *within* a given step (Hof et al., 2005), we contend that averaging *MoS*_*ML*_ values across steps does not usefully characterize a *distribution* (e.g., Fig. 2B) of step-to-step *MoS*_*ML*_ values. We assert that instead, one should calculate the statistical probability of a walker entering the Lateral “Instability” region (Fig. 2A) on any given step. Here, we propose the lateral Probability of Instability (*PoI*_*L*_) to compute just this.

Given a statistical distribution of *MoS*_*ML*_ (Eq. (1)) values (or equivalently, *δ*_*p*_ values; Fig. 2B), *PoI*_*L*_ is estimated by calculating the cumulative probability that values in that distribution will exhibit *MoS*_*ML*_ < 0. We thus integrate the probability density function of a given time series of *MoS*_*ML*_ over the range of *MoS*_*ML*_ < 0. Assuming a normally-distributed *MoS*_*ML*_ time series:

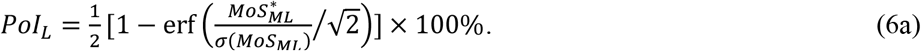

Equivalently, *PoI*_*L*_ can be computed from any *δ*_*P*_ time series as the probability that any *δ*_*P*_ will exceed the *MoS*_*ML*_ = 0 bound along *ê*_*P*_ (Fig 2B):

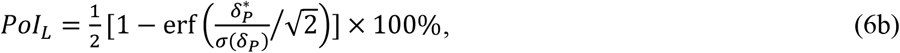

where 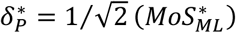 (Fig. 2A) and 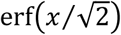 is the error function evaluated at *x* ((Abramowitz and Stegun, 1965); https://dlmf.nist.gov/). Because *PoI*_*L*_ incorporates the variance of *MoS*_*ML*_ (Eq. 6a) or mediolateral CoM state (Eq. 6b), it provides a measure of a participant’s lateral instability likelihood *on any given step* and helps explain shifts in mean *MoS*_*ML*_ for different conditions. This cannot be accomplished using average *MoS*_*ML*_ (Fig. 3). MATLAB code to compute *PoI* is available on GitHub (github.com/meghankazanski/PoI).

**Figure 3.**
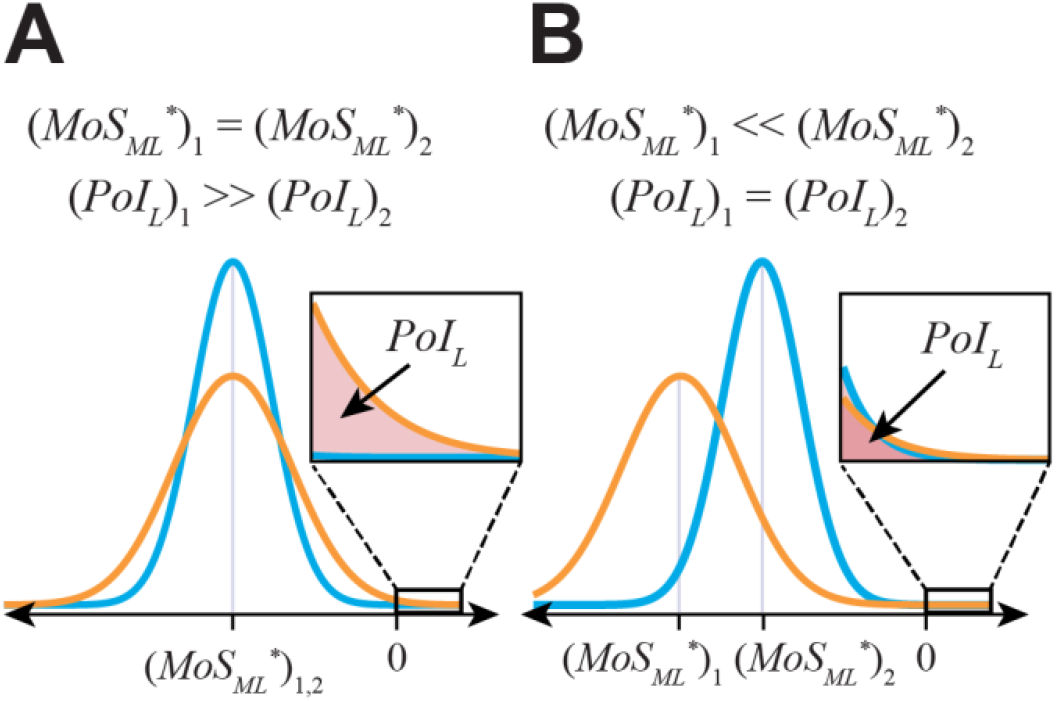
Probability of Instability (*PoI*_*L*_) depends on both the mean and variance of the distribution of any given *MoS*_*ML*_ time series. To demonstrate this, consider two hypothetical trial distributions in orange (1) and blue (2). In **(A)**, distributions with the exact same means 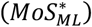 can still yield substantially different *PoI*_*L*_. Conversely, in **(B)**, distributions with very different means can still yield nearly identical *PoI*_*L*_.

We estimated *PoI*_*L*_ for each trial. We conducted a two-factor mixed-effects (Age×Condition) ANOVA (see *Supplement 2*) to assess age group and condition differences.

## RESULTS

All participants exhibited significantly larger *MoS*_*ML*_ means during perturbed walking (VIS and PLAT) relative to unperturbed (NOP) (Fig. 4A; p < 10^−3^; Table S2.1). All participants also demonstrated significantly more variable *MoS*_*ML*_ when perturbed (Fig. 4B; p < 10^−11^; Table S2.1). Participants generally exhibited slight anti-persistence of *MoS*_*ML*_ time series when perturbed (Fig. 4C; Table S2.1). YH adults exhibited significantly decreased DFA α during both VIS (p < 10^−2^) and PLAT (p < 10^−11^) perturbations. OH adults exhibited significantly decreased DFA α only during PLAT (p < 10^−9^) perturbations (Table S2.1). Age group effects were not statistically significant.

**Figure 4.**
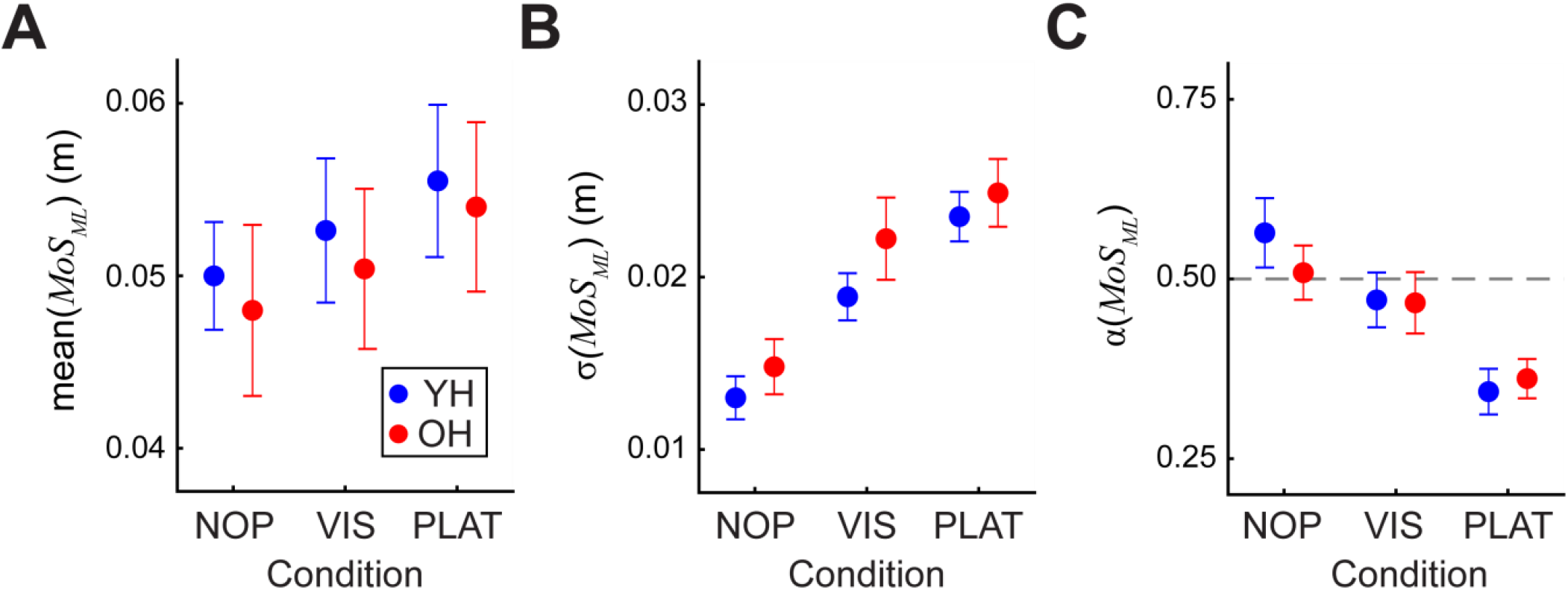
**(A)** *MoS*_*ML*_ mean, **(B)** variability (σ(*MoS*_*ML*_)) and **(C)** statistical persistence (DFA exponent: α(*MoS*_*ML*_)) for both YH and OH age groups for each of NOP, VIS and PLAT conditions. Error bars represent between-participant ±95% confidence intervals. During VIS and PLAT perturbations, participants demonstrated larger (p = 1.70×10^−15^), more-variable (p = 2.11×10^−65^), and increasingly anti-persistent (p = 1.68×10^−26^) *MoS*_*ML*,_ relative to the NOP condition. No statistically significant age group effects were observed (see *Supplement 2*; Table S2.1).

Across all conditions, step-to-step mediolateral CoM dynamics were strongly elongated along the candidate constant-stability GEM (Fig. 5). Under VIS and PLAT perturbations, distribution clouds of data points appeared larger relative to NOP (Fig. 5B-C), but remained strongly elongated along the constant-stability GEM.

**Figure 5.**
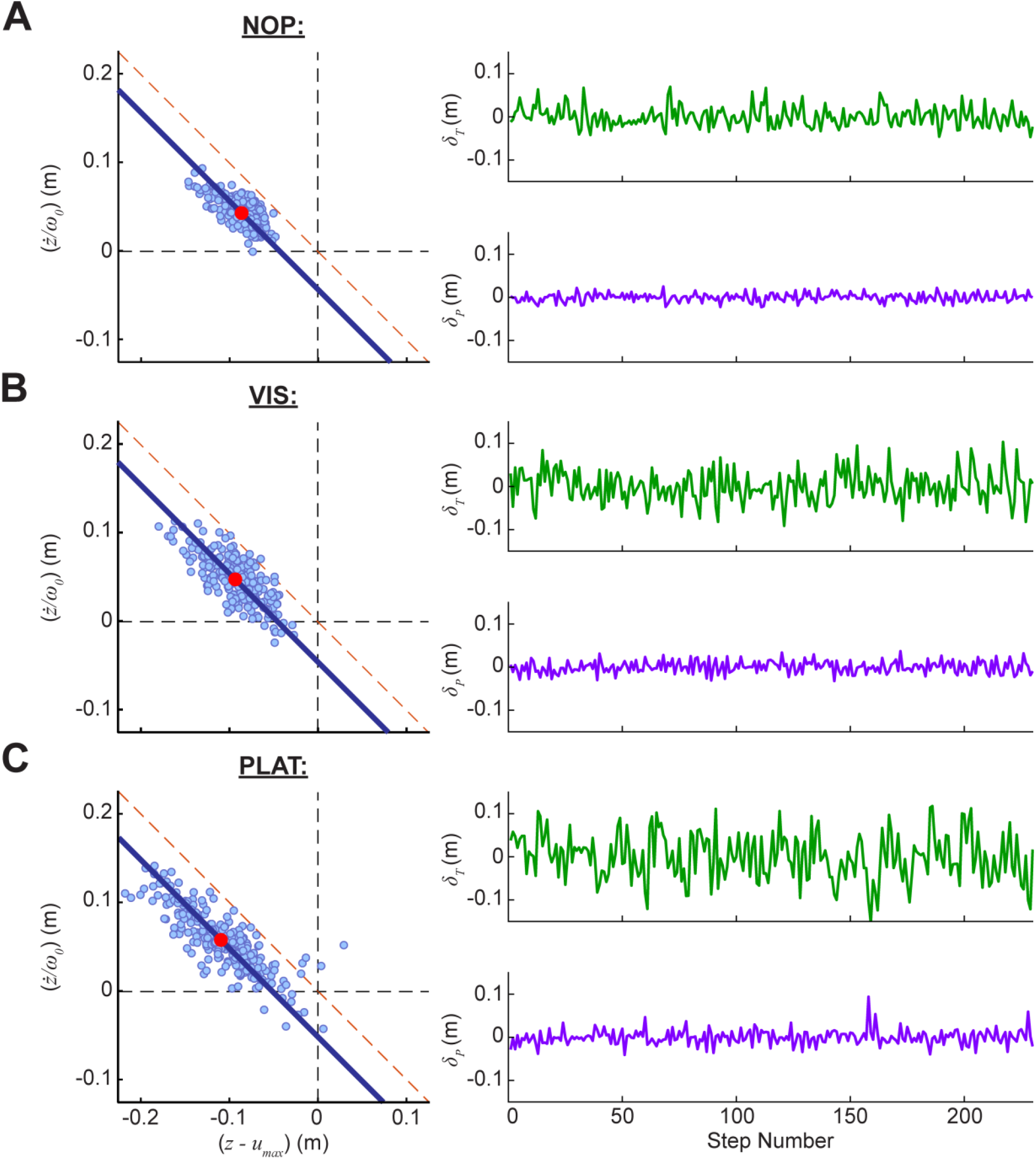
Example data from representative trials of a typical participant for each of (**A**) NOP, (**B**) VIS, and (**C**) PLAT conditions. **Left**: Mediolateral CoM dynamics attained at (*MoS*_*ML*_)_*n*_ for each *n*^th^ step (blue markers) of the corresponding trial. The candidate stability GEM (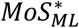 blue line) is defined by the trial-average: ⟨(*MoS*_*ML*_)_*n*_⟩_*N*_. The operating point ([(*z-u*_*max*_)^*\**^, (*ż/ω*_*0*_) ^*\**^]; red marker) on this manifold is likewise defined by the trial-average: [⟨(*z-u*_*max*_)_*n*_⟩_*N*_, ⟨(*ż/ω*_0_)_*n*_⟩_*N*_]. **Right**: Corresponding step-to-step time series of deviations from the operating point, tangent (*δ*_*T*_) and perpendicular (*δ*_*P*_) to the candidate stability GEM.

Across participants and conditions, *δ*_*P*_ deviations were significantly less variable than *δ*_*T*_ (Fig. 6A; σ(*δ*_*P*_) << σ(*δ*_*T*_); p = 1.15×10^−150^; Table S2.2). Likewise, σ(*δ*_*P*_) and σ(*δ*_*T*_) both increased significantly during perturbation conditions (Fig. 6A; all p < 10^−11^; Table S2.3). Across participants and conditions, *δ*_*P*_ fluctuations were either more statistically persistent (higher α > 0.5) or less anti-persistent (higher α < 0.5) (Fig. 6B; α(*δ*_*P*_) << α(*δ*_*T*_); p = 5.81×10^−16^; Table S2.2) than *δ*_*T*_ deviations. During perturbation conditions, both α(*δ*_*P*_) and α(*δ*_*T*_) generally became either less statistically persistent or even anti-persistent (Fig. 6B; Table S2.3). Age group effects were not significant.

**Figure 6.**
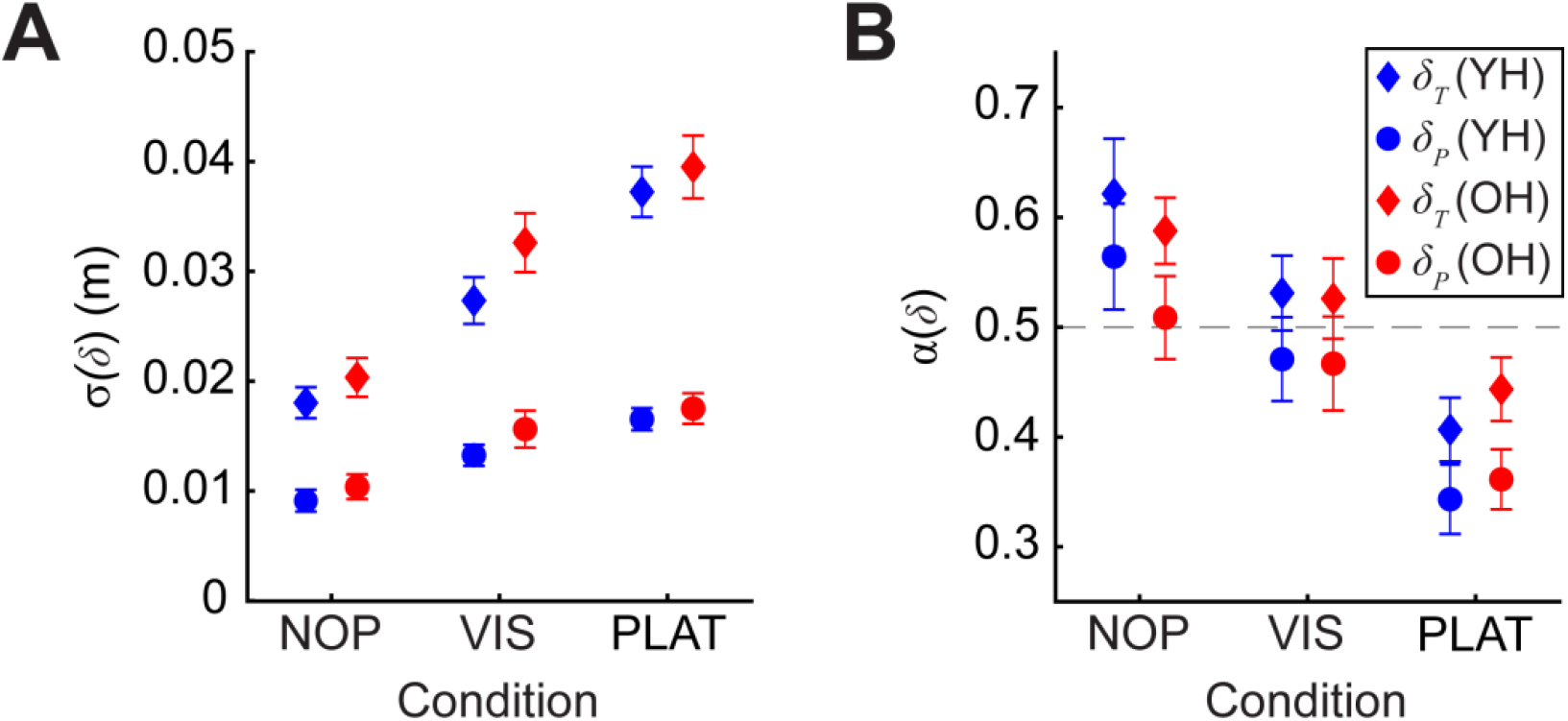
**(A):** Variability (σ(*δ*)) for all *δ*_*T*_ and *δ*_*P*_ deviation time series for YH and OH age groups, for each condition (NOP, VIS and PLAT). **(B):** DFA α exponents for all *δ*_*T*_ and *δ*_*P*_ time series for YH and OH age groups, for each condition. Error bars represent between-subject ± 95% confidence intervals. Goal-equivalent *δ*_*T*_ deviations were consistently more-variable (p = 1.15×10^−150^) and either more-persistent (higher α>0.5) or less anti-persistent (higher α<0.5) (p = 5.81×10^−16^) than goal-relevant *δ*_*P*_ deviations. During VIS and PLAT perturbations, participants demonstrated more-variable and either less-persistent or anti-persistent goal-equivalent *δ*_*T*_ (p = 1.35×10^−64^; p = 5.52×10^−22^) and goal-relevant *δ*_*P*_ (p = 2.11×10^−65^; p = 1.63×10^−26^) deviations, relative to NOP. No statistically significant age group effects were observed (see *Supplement 2*, Tables S2.2-3).

Across age groups and conditions, the extent to which participants regulated step width, α(*w*), and absolute lateral position, α(*z*_*B*_), strongly predicted the fluctuation dynamics, α(*δ*_*T*_) and α(*δ*_*P*_), of step-to-step CoM deviations (Fig. 7).

**Figure 7.**
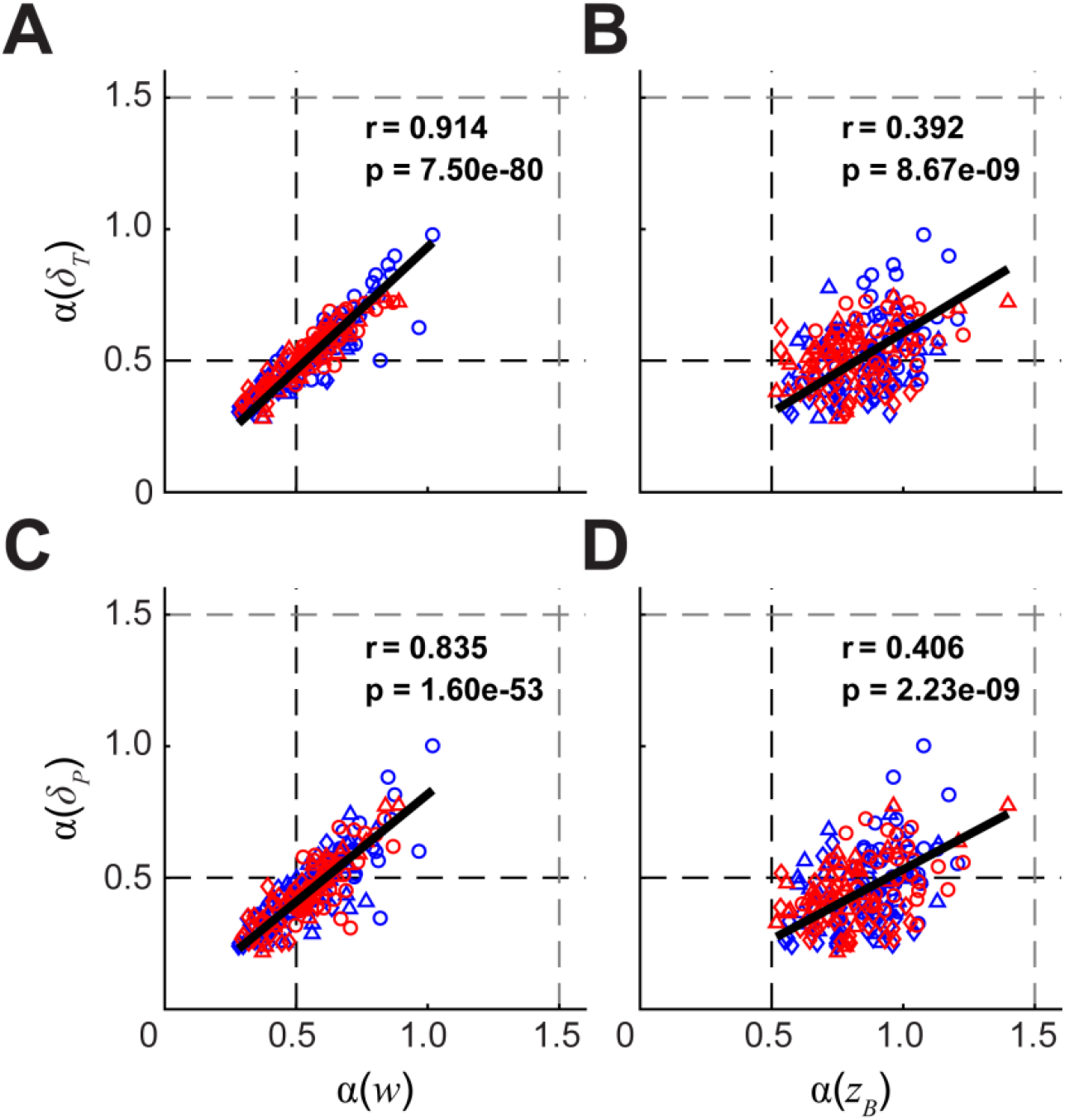
Pairwise Pearson correlations of DFA α exponents for time series of step width (*w*; **A** and **C**) and mediolateral body position (*z*_*B*_; **B** and **D**) with those of mediolateral CoM deviations tangent (*δ*_*T*_; **A** and **B**) and perpendicular (*δ*_*P*_; **C** and **D**) to the candidate stability GEM. Data were pooled across age groups and conditions. YH adults are visualized with blue markers, OH adults with red markers. Conditions are depicted as NOP (○), VIS (△), PLAT (◊). The step-to-step statistical persistence of both *w* and *z*_*B*_ stepping time series significantly predicted that of each of the *δ*_*T*_ and *δ*_*P*_ CoM time series.

Both YH and OH participants exhibited increased *PoI*_*L*_ (Fig. 8C; all p < 10^−9^; Table S2.4) when perturbed, indicating a *greater* likelihood of being laterally unstable on a given step, despite exhibiting larger average *MoS*_*ML*_ (Fig. 4A). This was due to increased *MoS*_*ML*_ (or *δ*_*P*_) variance (Fig. 8A-B). While OH demonstrated slightly larger *PoI*_*L*_, age group effects were not statistically significant.

**Figure 8.**
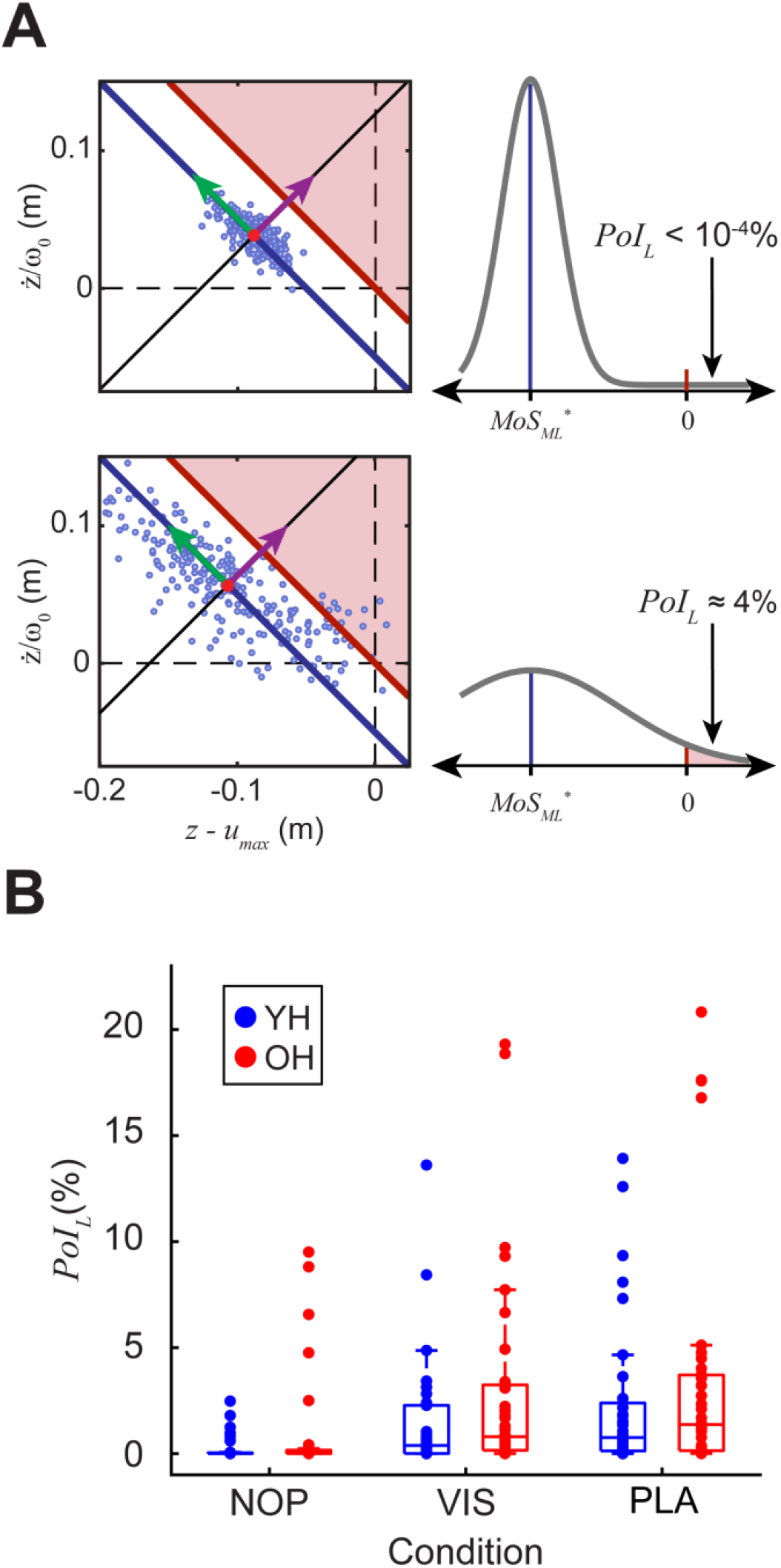
**(A):** Data from two representative trials (*top*: YH participant during NOP; *bottom*: OH participant during PLAT). Both participants demonstrated nearly identical average *MoS*_*ML*_ and hence nearly-identical candidate 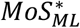 GEMs (*left*). However, the increased variance in the OH PLAT trial (*bottom*) produced substantially greater instability likelihood (*PoI*_*L*_; *right*). This confirms that average *MoS*_*ML*_ cannot capture instability likelihood (Fig. 3). **(B):** Box plots of *PoI*_*L*_ results for all trials for all participants. During VIS and PLAT perturbations, participants demonstrated significantly larger *PoI*_*L*_ (p = 9.45×10^−34^), relative to NOP. While some OH participants demonstrated larger *PoI*_*L*_, group differences were not statistically significant (p = 0.199) (see *Supplement 2*, Table S2.4).

## DISCUSSION

*MoS*_*ML*_ (Hof et al., 2005) has been widely used to assess lateral balance of human walking (Bruijn et al., 2013). However, when studies average *MoS*_*ML*_ across steps, a paradox arises. Numerous gait pathologies (e.g., (Watson et al., 2021), etc.) or destabilizing perturbations (e.g., (Onushko et al., 2019), etc.) induce walkers to exhibit larger (theoretically *more* stable) average *MoS*_*ML*_, despite clearly *de*stabilizing intrinsic and/or extrinsic factors. Multiple studies speculate that such counterintuitive findings might reflect compensatory mechanisms used to reduce likelihood of balance loss. However, this cannot explain why these individuals continue to exhibit elevated risk of balance loss despite such compensations. Moreover, averaging *MoS*_*ML*_ across steps discards all information pertaining to step-to-step dynamics that can provide insights into how people regulate variations across steps (Dingwell and Cusumano, 2019). Before considering whether *MoS*_*ML*_ is suitable to monitor balance impairments (Lencioni et al., 2021; Watson et al., 2021), much less address falling likelihood (Herssens et al., 2020), we must first resolve this paradox.

Here, we first demonstrated that balance dynamics in CoM state space (Bottaro et al., 2008; Hof, 2008) contains an infinite number of *constant-MoS*_*ML*_ *manifolds* (Fig. 2A) that arise directly from Hof’s definition of *MoS*_*ML*_ (Eqs. (1)-(2)). Importantly, these constant-*MoS*_*ML*_ manifolds exist regardless of how people might choose to regulate their stepping movements. Multiple studies, both computational (Bottaro et al., 2008; Hof, 2008) and experimental (Curtze et al., 2011; Rosenblatt and Grabiner, 2010), suggest people may try to maintain some constant 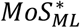. The GEM approach offers a unified framework to formally define a constant-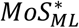 goal function (Eq. (4)) that allowed us to directly test this control hypothesis. Although *δ*_*P*_ deviations off of the constant-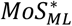 GEM were readily corrected (α(*δ*_*P*_) ≈ 0.5; Fig. 6B), humans did *not* exploit the equifinality *along* this candidate GEM. The *δ*_*T*_ deviations (α(*δ*_*T*_) ≈ 0.5; Fig. 6B) did not exhibit the statistical persistence (i.e., α >> 0.5) that would indicate they went *un*corrected, as is expected by the Minimal Intervention Principle (Todorov, 2004) because *δ*_*T*_ deviations have no bearing on *MoS*_*ML*_. Thus, we reject this hypothesis and conclude that defining a goal to simply maintain some constant 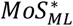 is insufficient to explain the step-to-step fluctuation dynamics observed in mediolateral CoM states (Figs. 5-6).

By its fundamental nature, steady walking is enacted by taking appropriate steps (Redfern and Schumann, 1994; Townsend, 1985): if you’re not taking steps, you’re not “walking” (Dingwell et al., 2010). However, one then needs to also regulate those stepping processes from step to step to be able to achieve *goal-directed*, or *meaningful* walking (Hof, 2008; Patil et al., 2022). Because walking humans can neither directly sense (Ting and Macpherson, 2004) nor directly actuate their CoM itself, it is plausible that they use other means like foot placement (Bruijn and van Dieën, 2018) to maintain balance. Our findings indeed demonstrated that the extent to which participants regulated their foot placements to maintain step width (*w*) (and, to a lesser extent, lateral position, *z*_*B*_) strongly predicted the fluctuation dynamics of both *δ*_*P*_ and *δ*_*T*_ deviations in mediolateral CoM states (Fig. 7). This supports the idea that humans *directly* regulate mediolateral foot placements (Dingwell and Cusumano, 2019; Kazanski et al., 2020) and that walking balance and CoM dynamics arise *in*directly as a consequence of this. Our findings are also consistent with neurophysiological evidence showing that distal sensory information underlies automatic postural responses for balance (Ting and Macpherson, 2004).

To address the problems inherent in interpreting average *MoS*_*ML*_, we introduced the lateral Probability of Instability (*PoI*_*L*_; Eqs. (6a)-(6b)) as an alternative metric that directly computes instability likelihood from step-to-step measurements of minimum *MoS*_*ML*_. Consistent with the aforementioned paradox, all participants here demonstrated larger average *MoS*_*ML*_ (Fig. 4A), despite being subjected to clearly *de*stabilizing mediolateral perturbations. Conversely, consistent with the nature of these perturbations, *PoI*_*L*_ (Fig. 8) revealed that participants’ increased variability (Figs. 4B & 6A) made them *more* likely to become laterally more unstable when perturbed (VIS and PLAT). Although no participants actually fell, they did become more unbalanced and thus needed to recruit active rebalancing mechanisms (Hof et al., 2005) to counteract these instances of lateral imbalance. *PoI*_*L*_ (Fig. 8B) correctly captured this increased risk of losing lateral balance, whereas average *MoS*_*ML*_ (Fig. 4A) clearly did not. Furthermore, by also incorporating variance of *MoS*_*ML*_, *PoI*_*L*_ explains why average *MoS*_*ML*_ increased in these destabilizing contexts: it was not because people became “more stable”, but because they moved their “stability operating point” (i.e., mean *MoS*_*ML*_) farther from the Hof stability boundary to accommodate increased variability.

As humans, we do not walk “on average” – we walk one step at a time. Likewise, we also do not fall or loose balance “on average”, but from specific events that occur in real time (Robinovitch et al., 2013). Our results explain why averaging *MoS*_*ML*_ measurements across steps yields the paradoxical findings often reported in the literature. Conversely, *PoI*_*L*_ properly accounts for the statistical distribution characteristics of *MoS*_*ML*_ across the “Stability” and “Lateral Instability” regions (Fig. 2B) to directly estimate single-step instability likelihood. Our unified framework thus re-frames Hof’s condition for dynamic stability (Hof et al., 2005) to incorporate intrinsic equifinality and step-to-step dynamics (Cusumano and Dingwell, 2013). From this, *PoI*_*L*_ emerges as a simple clinical statistic that resolves the *MoS*_*ML*_ paradox. While a substantial improvement, computing *PoI*_*L*_ does require time series of many steps and so may not be feasible in some clinical settings. Nevertheless, if the intent of applying *MoS*_*ML*_ is to assess or predict likelihood of imbalance (e.g., as in (Herssens et al., 2020; Lencioni et al., 2021; Watson et al., 2021)), then computing *PoI*_*L*_ instead of average *MoS*_*ML*_ will yield more meaningful conclusions regarding actual lateral instability risk.

## ACKNOWLEDGEMENTS

This work was supported by NIH grant 1-R01-AG049735.

## REFERENCES

Abramowitz, M., Stegun, I.A., 1965. Handbook of Mathematical Functions: with Formulas, Graphs, and Mathematical Tables. Dover Publications, Mineola, New York.

Bauby, C.E., Kuo, A.D., 2000. Active Control of Lateral Balance in Human Walking. J. Biomech. 33, 1433–1440.

Beltran, E.J., Dingwell, J.B., Wilken, J.M., 2014. Margins of stability in young adults with traumatic transtibial amputation walking in destabilizing environments. J. Biomech. 47, 1138–1143.

Bottaro, A., Yasutake, Y., Nomura, T., Casadio, M., Morasso, P., 2008. Bounded stability of the quiet standing posture: An intermittent control model. Hum. Mov. Sci. 27, 473–495.

Braun, D.A., Nagengast, A.J., Wolpert, D., 2011. Risk-sensitivity in sensorimotor control. Frontiers in Human Neuroscience 5.

Bruijn, S.M., Meijer, O.G., Beek, P.J., van Dieën, J.H., 2013. Assessing the stability of human locomotion: a review of current measures. J. R. Soc. Interface 10, 20120999.

Bruijn, S.M., van Dieën, J.H., 2018. Control of human gait stability through foot placement. J. R. Soc. Interface 15, 1–11.

Burns, E.R., Kakara, R., 2018. Deaths from Falls Among Persons Aged ≥65 Years — United States, 2007–2016. Morb. Mortal. Wkly. Rep. 67, 509–514.

Cho, H., Heijnen, M.J.H., Craig, B.A., Rietdyk, S., 2021. Falls in young adults: The effect of sex, physical activity, and prescription medications. PLoS ONE 16, e0250360.

Crenshaw, J.R., Bernhardt, K.A., Achenbach, S.J., Atkinson, E.J., Khosla, S., Kaufman, K.R., Amin, S.T., 2017. The circumstances, orientations, and impact locations of falls in community-dwelling older women. Archives of Gerontology and Geriatrics 73, 240–247.

Curtze, C., Hof, A.L., Postema, K., Otten, B., 2011. Over rough and smooth: Amputee gait on an irregular surface. Gait Posture 33, 292–296.

Cusumano, J.P., Cesari, P., 2006. Body-Goal Variability Mapping in an Aiming Task. Biol. Cybern. 94, 367–379.

Cusumano, J.P., Dingwell, J.B., 2013. Movement Variability Near Goal Equivalent Manifolds: Fluctuations, Control, and Model-Based Analysis. Hum. Mov. Sci. 32, 899–923.

Day, K.V., Kautz, S.A., Wu, S.S., Suter, S.P., Behrman, A.L., 2012. Foot placement variability as a walking balance mechanism post-spinal cord injury. Clin. Biomech. 27, 145–150.

Dean, J.C., Alexander, N.B., Kuo, A.D., 2007. The effect of lateral stabilization on walking in young and old adults. IEEE Trans. Biomed. Eng. 54, 1919–1926.

Dingwell, J.B., Cusumano, J.P., 2010. Re-Interpreting Detrended Fluctuation Analyses of Stride-To-Stride Variability in Human Walking. Gait Posture 32, 348–353.

Dingwell, J.B., Cusumano, J.P., 2019. Humans Use Multi-Objective Control to Regulate Lateral Foot Placement When Walking. PLoS Comput. Biol. 15, e1006850.

Dingwell, J.B., Cusumano, J.P., Rylander, J.H., Wilken, J.M., 2021. How Persons With Transtibial Amputation Regulate Lateral Stepping While Walking in Laterally Destabilizing Environments. Gait Posture 83, 88–95.

Dingwell, J.B., John, J., Cusumano, J.P., 2010. Do Humans Optimally Exploit Redundancy to Control Step Variability in Walking? PLoS Comput. Biol. 6, e1000856.

Gates, D.H., Scott, S.J., Wilken, J.M., Dingwell, J.B., 2013. Frontal plane dynamic margins of stability in individuals with and without transtibial amputation walking on a loose rock surface. Gait Posture 38, 570–575.

Havens, K.L., Mukherjee, T., Finley, J.M., 2018. Analysis of biases in dynamic margins of stability introduced by the use of simplified center of mass estimates during walking and turning. Gait Posture 59, 162–167.

Heijnen, M.J.H., Rietdyk, S., 2016. Falls in young adults: Perceived causes and environmental factors assessed with a daily online survey. Hum. Mov. Sci. 46, 86–95.

Herssens, N., van Criekinge, T., Saeys, W., Truijen, S., Vereeck, L., van Rompaey, V., Hallemans, A., 2020. An investigation of the spatio-temporal parameters of gait and margins of stability throughout adulthood. J. R. Soc. Interface 17, 20200194.

Hof, A.L., 2007. The equations of motion for a standing human reveal three mechanisms for balance. J. Biomech. 40, 451–457.

Hof, A.L., 2008. The ’extrapolated center of mass’ concept suggests a simple control of balance in walking. Hum. Mov. Sci. 27, 112–125.

Hof, A.L., Gazendam, M.G.J., Sinke, W.E., 2005. The condition for dynamic stability. J. Biomech. 38, 1–8.

Hof, A.L., van Bockel, R.M., Schoppen, T., Postema, K., 2007. Control of lateral balance in walking: Experimental findings in normal subjects and above-knee amputees. Gait Posture 25, 250–258.

Hsieh, K.L., Sheehan, R.C., Wilken, J.M., Dingwell, J.B., 2018. Healthy individuals are more maneuverable when walking slower while navigating a virtual obstacle course. Gait Posture 61, 466–472.

Hurt, C.P., Grabiner, M.D., 2015. Age-related differences in the maintenance of frontal plane dynamic stability while stepping to targets. J. Biomech. 48, 592–597.

John, J., Dingwell, J.B., Cusumano, J.P., 2016. Error Correction and the Structure of Inter-Trial Fluctuations in a Redundant Movement Task. PLoS Comput. Biol. 12, e1005118.

Kazanski, M.E., Cusumano, J.P., Dingwell, J.B., 2020. How Healthy Older Adults Regulate Lateral Stepping While Walking in Laterally Destabilizing Environments. J. Biomech. 104, 109714.

Lencioni, T., Anastasi, D., Carpinella, I., Castagna, A., Crippa, A., Gervasoni, E., Marzegan, A., Rabuffetti, M., Pelosin, E., Cattaneo, D., Ferrarin, M., 2021. Strategies for maintaining dynamic balance in persons with neurological disorders during overground walking. Proceedings of the Institution of Mechanical Engineers, Part H: Journal of Engineering in Medicine 235, 1079–1087.

Li, J., Huang, H.J., 2022. Small directional treadmill perturbations induce differential gait stability adaptation. J. Neurophysiol. 127, 38–55.

Maraun, D., Rust, H.W., Timmer, J., 2004. Tempting long-memory - on the interpretation of DFA results. Nonlinear Processes in Geophysics 11, 495–503.

McAndrew Young, P.M., Wilken, J.M., Dingwell, J.B., 2012. Dynamic Margins of Stability During Human Walking in Destabilizing Environments. J. Biomech. 45, 1053–1059.

Nagengast, A.J., Braun, D.A., Wolpert, D.M., 2011. Risk-sensitivity and the mean-variance trade-off: decision making in sensorimotor control. Proc. Roy. Soc. B Biol. Sci. 278, 2325–2332.

Onushko, T., Boerger, T., Van Dehy, J., Schmit, B.D., 2019. Dynamic stability and stepping strategies of young healthy adults walking on an oscillating treadmill. PLoS ONE 14, e0212207.

Patil, N.S., Dingwell, J., Cusumano, J.P., 2019. Correlations of pelvis state to foot placement do not imply within-step active control. J. Biomech. 97, 109375.

Patil, N.S., Dingwell, J.B., Cusumano, J.P., 2022. Viability, Task Switching, and Fall Avoidance of the Simplest Dynamic Walker. Scientific Reports 12, 8993.

Peebles, A.T., Reinholdt, A., Bruetsch, A.P., Lynch, S.G., Huisinga, J.M., 2016. Dynamic margin of stability during gait is altered in persons with multiple sclerosis. J. Biomech. 49, 3949–3955.

Redfern, M.S., Schumann, T., 1994. A model of foot placement during gait. J. Biomech. 27, 1339–1346.

Render, A.C., Kazanski, M.E., Cusumano, J.P., Dingwell, J.B., 2021. Walking humans trade off different task goals to regulate lateral stepping. J. Biomech. 119, 110314.

Rethwilm, R., Böhm, H., Haase, M., Perchthaler, D., Dussa, C.U., Federolf, P., 2021. Dynamic stability in cerebral palsy during walking and running: Predictors and regulation strategies. Gait Posture 84, 329–334.

Robinovitch, S.N., Feldman, F., Yang, Y., Schonnop, R., Leung, P.M., Sarraf, T., Sims-Gould, J., Loughin, M., 2013. Video capture of the circumstances of falls in elderly people residing in long-term care: an observational study. Lancet 381, 47–54.

Rodrigues, F.B., deSáe Souza, G.S., de Mendonça Mesquita E., de Sousa Gomide, R., Baptista, R.R., Pereira, A.A., Andrade, A.O., Vieira, M.F., 2021. Margins of stability of persons with transtibial or transfemoral amputations walking on sloped surfaces. J. Biomech. 123, 110453.

Rosenblatt, N.J., Grabiner, M.D., 2010. Measures of frontal plane stability during treadmill and overground walking. Gait Posture 31, 380–384.

Ting, L.H., Macpherson, J.M., 2004. Ratio of Shear to Load Ground-Reaction Force May Underlie the Directional Tuning of the Automatic Postural Response to Rotation and Translation. J. Neurophysiol. 92, 808–823.

Tisserand, R., Armand, S., Allali, G., Schnider, A., Baillieul, S., 2018. Cognitive-motor dual-task interference modulates mediolateral dynamic stability during gait in post-stroke individuals. Hum. Mov. Sci. 58, 175–184.

Todorov, E., 2004. Optimality principles in sensorimotor control. Nat. Neurosci. 7, 907–915.

Townsend, M.A., 1985. Biped Gait Stabilization Via Foot Placement. J. Biomech. 18, 21–38.

Wang, M., Wu, F., Callisaya, M.L., Jones, G., Winzenberg, T., 2021. Incidence and circumstances of falls among middle-aged women: a cohort study. Osteoporosis International 32, 505–513.

Wang, Y., Srinivasan, M., 2014. Stepping in the direction of the fall: the next foot placement can be predicted from current upper body state in steady-state walking. Biol. Lett. 10, 20140405.

Watson, F., Fino, P.C., Thornton, M., Heracleous, C., Loureiro, R., Leong, J.J.H., 2021. Use of the margin of stability to quantify stability in pathologic gait – a qualitative systematic review. BMC Musculoskeletal Disorders 22, 597.

Yang, Y., Komisar, V., Shishov, N., Lo, B., Korall, A.M.B., Feldman, F., Robinovitch, S.N., 2020. The Effect of Fall Biomechanics on Risk for Hip Fracture in Older Adults: A Cohort Study of Video-Captured Falls in Long-Term Care. Journal of Bone and Mineral Research 35, 1914–1922.

Zeni, J.A., Richards, J.G., Higginson, J.S., 2008. Two simple methods for determining gait events during treadmill and overground walking using kinematic data. Gait Posture 27, 710–714.

