## Supplement 1 for "Rethinking Margin of Stability: Incorporating Step-To-Step Regulation to Resolve the Paradox"

***Journal of Biomechanics***

### SUPPLEMENT 1: ADDITIONAL EXPERIMENTAL METHODS

This supplement provides additional details regarding participants, preliminary assessments, the experimental protocol, as well as kinematic data collection and processing methods pertaining to the main manuscript. Complete descriptions are also presented in Kazanski et al. (2020). We conducted the analyses presented here on data from the same cohort of young healthy (YH) and older healthy (OH) adults.

#### *Participants*

Seventeen young healthy (YH) and 19 older healthy (OH) adults participated in the study. All participants provided written informed consent, as approved by The University of Texas at Austin's IRB, and had no physiological conditions or limitations that affected their walking. All participants were required to score ≥24/30 on the Mini-Mental State Exam (Folstein et al., 1975). Two OH participants relied heavily on the handrails during walking trials. These data were not included in any analyses. We conducted all analyses on the remaining 17 YH and 17 OH adults.

We measured each participant's body mass, height, leg length, and body mass index. We assessed each participant's general physical capacity by evaluating functional mobility (Timed Up and Go (Podsiadlo and Richardson, 1991)), dynamic stepping capability (Four Square Step Test (Dite and Temple, 2002)), average maximum left/right isometric quadriceps strength (Salinas et al., 2017), and preferred overground walking speed. We assessed each participant's falling concern using the 10-point Iconographic-Falls Efficacy Scale (Delbaere et al., 2011). Two-sample t-tests evaluated YH vs. OH differences in physical characteristics and assessment scores (Table S1.1).

**Table S1.1** – Additional participant characteristics and assessment scores for Young (YH) and Older (OH) healthy adults (see also Table 1 in main manuscript). All values except Sex are given as mean ± standard deviation. Two-sample t-test results (p-values) for Age group differences are shown.

| **Characteristic:** | **Young Healthy (YH):** | **Older Healthy (OH):** | **p-value** |
|---|---|---|---|
| Sex [M/F] | 8 / 9 | 7 / 10 | N/A |
| Body Mass [kg] | 64.5 ± 12.5 | 73.3 ± 18.9 | 0.118 |
| Body Mass Index [kg/m$^2$] | 21.7 ± 3.2 | 24.9 ± 5.4 | **0.042** |
| **Assessment:** | | | |
| Timed Up and Go [s] | 8.06 ± 1.20 | 8.66 ± 1.08 | 0.139 |
| Four Square Step Test [s] | 7.18 ± 1.53 | 9.07 ± 2.07 | **0.005** |
| Preferred Walking Speed [m/s] | 1.46 ± 0.15 | 1.35 ± 0.14 | **0.033** |

#### *Experimental Protocol*

Participants walked on an instrumented treadmill in a "V-Gait" virtual reality system (Motekforce Link, Amsterdam, Netherlands) (Fig. S1.1). For each participant, treadmill and visual optic flow speeds were fixed at 90% of their preferred overground walking speed.

Following a 3-minute acclimation trial, participants completed three, 3-minute trials of treadmill walking for each of 3 conditions: normal walking with no perturbations (NOP), or walking with mediolateral oscillations of either the visual field (VIS) or treadmill platform (PLAT). All oscillations were confined to the mediolateral ($z$) direction (O'Connor and Kuo, 2009). These oscillations were continuous and pseudo-randomly administered to prevent entrainment (Warren et al., 1996), applied as a composite of sines with several incommensurate frequencies (similar to (McAndrew Young et al., 2012)):

$$D(t) = A \cdot [1.0 \sin(0.16 \cdot 2\pi t) + 0.8 \sin(0.21 \cdot 2\pi t) + 1.4 \sin(0.24 \cdot 2\pi t) + 0.5 \sin(0.49 \cdot 2\pi t)], \quad (1)$$

where *D(t)* is the mediolateral translation distance (m), *A* is a scaling factor, and *t* is time (s). During VIS trials (*A* = 0.25), the visual field translated according to Eq. (1), while the treadmill remained stationary. During PLAT trials (*A* = 0.015), the treadmill platform translated according to Eq. (1), while visual optic flow was normal. The virtual scene was a forest path lined by white posts (2.4 m tall; every 3 m), which were included to increase motion parallax (Bardy et al., 1996; McAndrew et al., 2010).

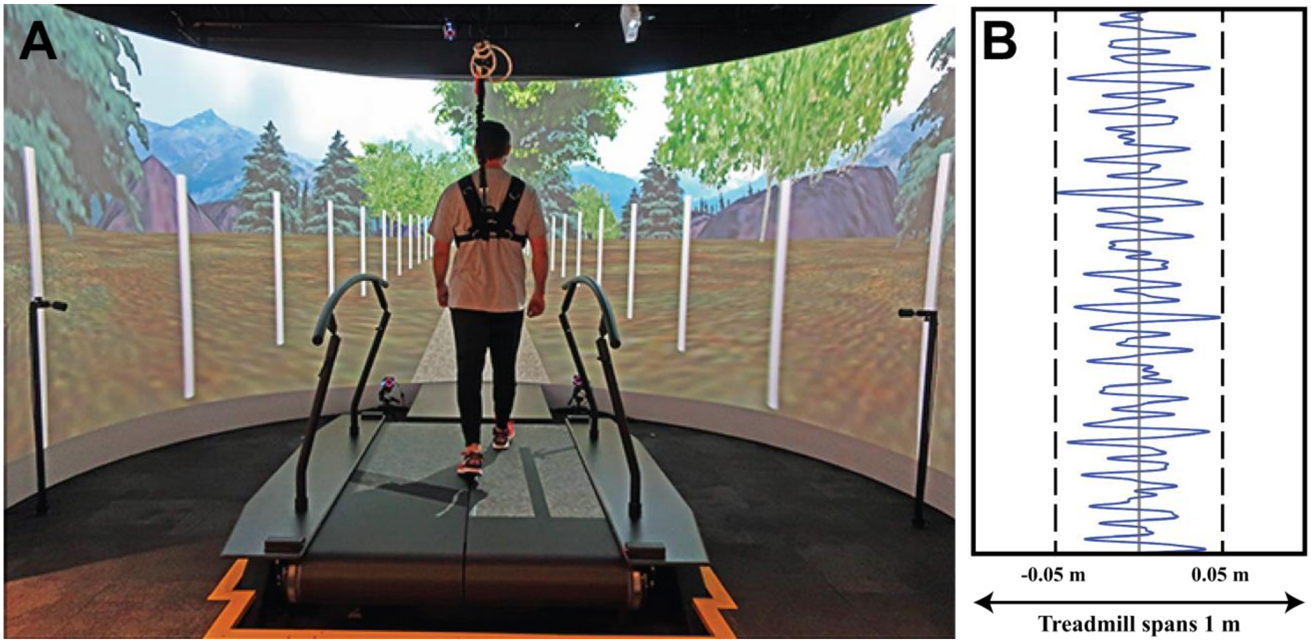


**Figure S1.1 A)** A typical participant walking on an instrumented treadmill along a virtual forest path lined with white posts. **B)** Sample sequence of the continuous, pseudorandom oscillations of the treadmill platform, which were confined to the mediolateral direction. Visual field oscillations (not shown) applied a similar sequence. This figure is adapted from Kazanski et al. (2020).

The order of condition presentation was balanced across participants. All same-condition trials were presented consecutively. The first trial of each perturbation condition (VIS and PLAT) acclimated participants by gradually increasing perturbation magnitudes from 50-100% during the first 2 minutes of the trial. Participants rested (≥ 2 min) between each walking trial.

*Kinematic Data Collection and Processing*

Each subject wore 16 retroreflective markers: four on the head (left and right forehead, left and right back head), four on the pelvis (left and right Posterior Superior Iliac Spine (PSIS), left and right Anterior Superior Iliac Spine (ASIS)), and four on each left and right shoe surface, aligned with various foot landmarks (lateral malleolus, calcaneus, first and fifth metatarsal heads). Four additional markers were placed on the treadmill platform to allow us to capture the kinematics of the treadmill reference frame, which oscillated during PLAT trials. Kinematic data were collected at 120 Hz using a 10-camera Vicon motion capture system (Oxford Metrics, Oxford, UK). Raw data were processed using Nexus (Oxford Metrics, Oxford, UK) and kinematic marker trajectories were exported to MATLAB (MathWorks, Natick, MA) for all further analyses.

Three PLAT trials (2 YH, 1 OH) were not analyzed due to excessive marker dropout. Marker trajectory data were filtered ($4^{th}$-order, low-pass Butterworth; cutoff = 6 Hz) and interpolated to 600 Hz to ensure accurate stepping event detection (Bohnsack-McLagan et al., 2016). Heel-strike and toe-off events were defined for each step (Zeni et al., 2008). We analyzed the $N$=230 consecutive steps following the first 15s of each trial.

---

### REFERENCES


Bardy, B.G., Warren, W.H., Kay, B.A., 1996. Motion Parallax is Used to Control Postural Sway During Walking. Experimental Brain Research 111, 271-282.

Bohnsack-McLagan, N.K., Cusumano, J.P., Dingwell, J.B., 2016. Adaptability of Stride-To-Stride Control of Stepping Movements in Human Walking. Journal of Biomechanics 49, 229-237.

Delbaere, K., T. Smith, S., Lord, S.R., 2011. Development and Initial Validation of the Iconographical Falls Efficacy Scale. The Journals of Gerontology Series A: Biological Sciences and Medical Sciences 66A, 674-680.

Dite, W., Temple, V.A., 2002. A clinical test of stepping and change of direction to identify multiple falling older adults. Archives of Physical Medicine and Rehabilitation 83, 1566-1571.

Folstein, M.F., Folstein, S.E., McHugh, P.R., 1975. “Mini-mental state”: A practical method for grading the cognitive state of patients for the clinician. Journal of Psychiatric Research 12, 189-198.

Kazanski, M.E., Dingwell, J.B., Cusumano, J.P., 2020. How healthy older adults regulate lateral foot placement while walking in laterally destabilizing environments. Journal of Biomechanics 104, 109714.

McAndrew, P.M., Dingwell, J.B., Wilken, J.M., 2010. Walking variability during continuous pseudo-random oscillations of the support surface and visual field. Journal of Biomechanics 43, 1470-1475.

McAndrew Young, P.M., Wilken, J.M., Dingwell, J.B., 2012. Dynamic margins of stability during human walking in destabilizing environments. Journal of Biomechanics 45, 1053-1059.

O'Connor, S.M., Kuo, A.D., 2009. Direction-Dependent Control of Balance During Walking and Standing. Journal of Neurophysiology 102, 1411-1419.

Podsiadlo, D., Richardson, S., 1991. The Timed Up and Go Test - A Test of Basic Functional Mobility for Frail Elderly Persons. Journal of the American Geriatrics Society 39, 142-148.

Salinas, M.M., Wilken, J.M., Dingwell, J.B., 2017. How Humans Use Visual Optic Flow to Regulate Stepping During Walking. Gait & Posture 57, 15-20.

Warren, W.H., Kay, B.A., Yilmaz, E.H., 1996. Visual Control of Posture During Walking: Functional Specificity. Journal of Experimental Psychology: Human Perception and Performance 22, 818-838.

Zeni, J.A., Richards, J.G., Higginson, J.S., 2008. Two simple methods for determining gait events during treadmill and overground walking using kinematic data. Gait & Posture 27, 710-714.
