## Supplement 2 for "Rethinking Margin of Stability: Incorporating Step-To-Step Regulation to Resolve the Paradox"

***Journal of Biomechanics***

### SUPPLEMENT 2: STATISTICAL METHODS & RESULTS

This supplement provides additional methodological details and outcomes of statistical analyses pertaining to the main manuscript. Analyses were performed in Minitab (Minitab, Inc. State College, PA), with significance determined at $p < 0.05$.

*Mediolateral Margin of Stability ($MoS_{ML}$)*

We applied a two-factor (Age×Condition) analysis of variance (ANOVA) using a mixed effects model to each of $MoS_{ML}$ means, variability, and DFA-α data sets. These models evaluated for Age (YH vs. OH) and Condition (NOP, VIS, and PLAT) main effects, and Age×Condition interaction effects. Standard deviations and DFA-α exponents were first log-transformed to fulfill ANOVA linearity and normality assumptions. When Condition main effects were significant, Tukey's post-hoc least significant difference pairwise comparisons were conducted for the relevant Condition differences across both Age groups. When Age×Condition interaction effects were significant, Tukey's post-hoc comparisons were conducted separately to assess Age group differences (YH vs. OH) for each Condition (NOP, VIS, and PLAT), as well as the relevant Condition differences (VIS vs. NOP and PLAT vs. NOP) for each Age group (YH and OH). Statistical results are presented in Table S1.

**Table S2.1** – Statistical results for differences between Age groups (YH vs. OH) and Conditions (NOP, VIS and PLAT) for the data shown in Fig. 4: $MoS_{ML}$ means, variability ($\sigma(MoS_{ML})$) and DFA exponents ($\alpha(MoS_{ML})$). ANOVA results (F-statistics and p-values) are provided for main effects of Age, Condition, and Age×Condition interaction effects, with relevant Tukey's post-hoc comparisons. Age group differences for each condition were all non-significant (all p>0.3). Significant differences are indicated in bold.

| Fig. | Variable | Age | Condition | Age × Condition | Tukey's (Condition) | Tukey's (Age×Condition) |
|---|---|---|---|---|---|---|
| 4A | Mean ($MoS_{ML}$) | $F_{(1,32)}=0.25$<br>$p = 0.168$ | **$F_{(2,163)} = 42.19$**<br>**$p = 1.70\times10^{-15}$** | $F_{(2,163)} = 0.03$<br>$p = 0.975$ | **PLAT-NOP: $p < 10^{-11}$**<br>**VIS-NOP: $p < 10^{-3}$** | N/A |
| 4B | Ln [$\sigma(MoS_{ML})$] | $F_{(1,32)}=1.66$<br>$p = 0.206$ | **$F_{(2,163)} = 425.16$**<br>**$p = 2.11\times10^{-65}$** | **$F_{(2,163)} = 3.90$**<br>**$p = 0.022$** | N/A | YH:<br>**PLAT-NOP: $p < 10^{-11}$**<br>**VIS-NOP: $p < 10^{-11}$**<br>OH:<br>**PLAT-NOP: $p < 10^{-11}$**<br>**VIS-NOP: $p < 10^{-11}$** |
| 4C | Ln [$\alpha(MoS_{ML})$] | $F_{(1,32)}=0.13$<br>$p = 0.725$ | **$F_{(2,163)} = 87.60$**<br>**$p = 1.68\times10^{-26}$** | **$F_{(2,163)} = 3.48$**<br>**$p = 0.033$** | N/A | YH:<br>**PLAT-NOP: $p < 10^{-11}$**<br>**VIS-NOP: $p < 10^{-2}$**<br>OH:<br>**PLAT-NOP: $p < 10^{-9}$**<br>VIS-NOP: $p =0.379$ |

*Candidate Stability GEM-Aligned Deviations ($\delta_T$ , $\delta_P$)*

To first test for Direction differences, we applied a three-factor (Age × Condition × Direction) analysis of variance (ANOVA) using a mixed effects model to $\delta_T$ and $\delta_P$ variability and DFA-α data sets. These models evaluated potential Age (YH vs. OH), Condition (NOP, VIS, and PLAT) and Direction ($\hat{e}_T$ vs. $\hat{e}_P$) main effects. Standard deviations and DFA-α exponents were first log-transformed to fulfill ANOVA linearity and normality assumptions. Statistical results are presented in Table S2.

**Table S2.2** – Statistical results for differences between Age groups (YH vs. OH), Conditions (NOP, VIS and PLAT) and Candidate Stability GEM-aligned deviation directions ($\hat{e}_T$ vs. $\hat{e}_P$) for the data shown in Fig. 6: $\delta_T$ and $\delta_P$ variability ($\sigma(\delta)$) and DFA exponents ($\alpha(\delta)$). ANOVA results (F-statistics and p-values) are provided for main effects of Age, Condition, and Direction. Significant differences are indicated in bold.

| Fig. | Variable | Age | Condition | Direction |
|---|---|---|---|---|
| 6A | Ln [σ(δ)] | $F_{(1,32)}$= 2.64<br>p = 0.114 | **$F_{(2,358)}$ = 522.14<br>p = 7.27×10$^{-107}$** | **$F_{(1,358)}$ = 2063.30<br>p = 1.15×10$^{-150}$** |
| 6B | Ln [α(δ)] | $F_{(1,32)}$=0.02<br>p = 0.893 | **$F_{(2,358)}$ = 155.44<br>p = 4.37×10$^{-49}$** | **$F_{(1,358)}$ = 72.00<br>p = 5.81×10$^{-16}$** |

Then, to test for Age group and walking Condition effects on GEM-aligned deviations in each direction, we applied a two-factor (Age×Condition) analysis of variance (ANOVA) using a mixed effects model to $\delta_T$ and $\delta_P$ variability and DFA-α data sets. These models evaluated for Age (YH vs. OH) and Condition (NOP, VIS, and PLAT) main effects, and Age×Condition interaction effects separately for each of σ($\delta_T$), σ($\delta_P$), α($\delta_T$), and α($\delta_P$) data sets. All data sets were first log-transformed to fulfill ANOVA linearity and normality assumptions. When Condition main effects were significant, we conducted Tukey's post-hoc least significant difference pairwise comparisons for the relevant Condition differences across both Age groups (YH and OH). When Age×Condition interaction effects were significant, we conducted Tukey's post-hoc comparisons separately to assess Age group differences (YH vs. OH) for each Condition (NOP, VIS, and PLAT), as well as the relevant Condition differences (VIS vs. NOP and PLAT vs. NOP) for each Age group (YH and OH). Statistical results are presented in Table S3.

*Lateral Probability of Instability ($PoI_L$)*

We applied a two-factor (Age×Condition) analysis of variance (ANOVA) using a mixed effects model to the $PoI_L$ data set. These models evaluated for Age (YH vs. OH) and Condition (NOP, VIS, and PLAT) main effects, and Age×Condition interaction effects. $PoI_L$ were first log-transformed to fulfill ANOVA linearity and normality assumptions. When Condition main effects were significant, we conducted Tukey's post-hoc comparisons for relevant Condition differences across both Age groups (YH and OH). When Age×Condition interaction effects were significant, we conducted Tukey's post-hoc comparisons separately to assess Age group differences (YH vs. OH) for each Condition (NOP, VIS, and PLAT), as well as the relevant Condition differences (VIS vs. NOP and PLAT vs. NOP) for each Age group (YH and OH). Statistical results are presented in Table S4.

**Table S2.3** – Statistical results for differences between Age groups (YH vs. OH) and Conditions (NOP, VIS and PLAT) for the data shown in Fig. 6: for each of $\delta_T$ and $\delta_P$, the variability ($\sigma(\delta_T)$ and $\sigma(\delta_P)$) and DFA exponents ($\alpha(\delta_T)$ and $\alpha(\delta_P)$). ANOVA results (F-statistics and p-values) are provided for main effects of Age, Condition, and Age×Condition interaction effects, with relevant Tukey's post-hoc comparisons. Age group differences for each condition were all non-significant (all p>0.5). Significant differences are indicated in bold.

| Fig. | Variable | Age | Condition | Age × Condition | Tukey's (Condition) | Tukey's (Age×Condition) |
|---|---|---|---|---|---|---|
| 6A | Ln [$\sigma(\delta_T)$] | $F_{(1,32)} = 2.44$<br>$p = 0.128$ | **$F_{(2,163)} = 413.75$**<br>**$p = 1.35\times10^{-64}$** | $F_{(2,163)} = 3.05$<br>$p = 0.050$ | **PLAT-NOP: $p < 10^{-11}$**<br>**VIS-NOP: $p < 10^{-11}$** | N/A |
| 6A | Ln [$\sigma(\delta_P)$] | $F_{(1,32)} = 1.66$<br>$p = 0.206$ | **$F_{(2,163)} = 425.15$**<br>**$p = 2.11\times10^{-65}$** | **$F_{(2,163)} = 3.90$**<br>**$p = 0.022$** | N/A | YH:<br>**PLAT-NOP: $p < 10^{-11}$**<br>**VIS-NOP: $p < 10^{-11}$**<br>OH:<br>**PLAT-NOP: $p < 10^{-11}$**<br>**VIS-NOP: $p < 10^{-11}$** |
| 6B | Ln [$\alpha(\delta_T)$] | $F_{(1,32)} = 0.04$<br>$p = 0.835$ | **$F_{(2,163)} = 67.07$**<br>**$p = 5.52\times10^{-22}$** | $F_{(2,163)} = 2.52$<br>$p = 0.084$ | **PLAT-NOP: $p < 10^{-10}$**<br>**VIS-NOP: $p < 10^{-4}$** | N/A |
| 6B | Ln [$\alpha(\delta_P)$] | $F_{(1,32)} = 0.13$<br>$p = 0.725$ | **$F_{(2,163)} = 87.60$**<br>**$p = 1.63\times10^{-26}$** | **$F_{(2,163)} = 3.48$**<br>**$p = 0.033$** | N/A | YH:<br>**PLAT-NOP: $p < 10^{-11}$**<br>**VIS-NOP: $p < 10^{-2}$**<br>OH:<br>**PLAT-NOP: $p < 10^{-9}$**<br>VIS-NOP: $p = 0.379$ |

**Table S2.4** – Statistical results for differences between Age groups (YH vs. OH) and Conditions (NOP, VIS and PLAT) for the data shown in Fig. 8C: the lateral Probability of Instability ($PoI_L$). ANOVA results (F-statistics and p-values) are provided for main effects of Age, Condition, and Age×Condition interaction effects, with relevant Tukey's post-hoc comparisons. Age group differences for each condition were all non-significant (all p>0.3). Significant differences are indicated in bold.

| Fig. | Variable | Age | Condition | Age × Condition | Tukey's (Condition) | Tukey's (Age×Condition) |
|---|---|---|---|---|---|---|
| 8C | Ln[$PoI_L$] | $F_{(1,32)} = 1.72$<br>$p = 0.199$ | **$F_{(2,163)} = 125.69$**<br>**$p = 9.45\times10^{-34}$** | **$F_{(2,163)} = 5.24$**<br>**$p = 0.006$** | N/A | YH:<br>**PLAT-NOP: $p < 10^{-11}$**<br>**VIS-NOP: $p < 10^{-11}$**<br>OH:<br>**PLAT-NOP: $p < 10^{-9}$**<br>**VIS-NOP: $p < 10^{-11}$** |
